# A Two-Filter Adaptation to Achieve Hemodiafiltration with Enhanced Performance

**DOI:** 10.1101/2023.06.16.545389

**Authors:** Kyle Chu, Pei Li, Irfani Ausri, Cesar Vasconez, Xiaowu (Shirley) Tang

**Author notes:** **Corresponding Author:** X. S. Tang, Department of Chemistry & Waterloo Institute for Nanotechnology, University of Waterloo, 200 University Ave West, Waterloo, Ontario, Canada N2L 3G1.

## Abstract

**Introduction:** Advancements in hemodialysis (HD) instrumentation have resulted in numerous breakthroughs in technology and significantly enhanced patient outcomes. Today, hemodiafiltration (HDF) which combines HD and hemofiltration has been widely used as an alternative to conventional HD in many countries. HDF is known to outperform conventional HD, offering more effective waste clearance and better fluid balance to patients. However, HDF requires newer-generation machines that are not accessible in many under-resourced geographical regions and societies. This study investigates a facile adaptation of conventional HD machines to achieve HDF. The objectives are to address the premature obsolescence of older but fully functional machines and to advocate for equal access to improved medical care and treatment.

**Methods:** A bench-top experimental setup was established to evaluate the performance of HDF using a two-filter adaptation in comparison to that of standard HD. Urea clearance, human serum album loss, and hemolysis were assessed under identical operational conditions for both configurations.

**Findings:** Our results show that the HDF configuration outperformed the HD configuration, with significantly higher urea clearance (268.31±44.17 mL/min via HDF vs. 53.33±13.20 mL/min via HD), but comparable human serum albumin loss and hemolysis levels.

**Discussion:** The explored two-filter adaptation presents a cost-effective method to achieve HDF with improved performance using conventional HD machines, with no added risk to patients. Further validation on patients in a hospital setting is necessary.

## INTRODUCTION

The performance of hemodialysis machines has greatly improved over the past few decades, with newer machines offering more precise flow control from simple extracorporeal circulation devices without volumetric control, with low blood flow and low-flux dialyzers to precisely controlling devices.^[1,2]^ At an ever-increasing rate, the efficiency of older machines becomes less desirable, which is highly related to the performance of dialysis. However, hemodialysis machines are costly pieces of equipment. In under-resourced geographical regions and societies, purchasing a new set of machines at each dialysis clinic is not always economically viable or environmentally responsible. As such, cost-effective methods to alter older but fully functional machines to achieve performance comparable to that of newer machines are highly desirable. This approach aligns with the United Nations’ objective of promoting equitable access to healthcare, which seeks to ensure that all individuals have access to high-quality healthcare services regardless of their economic or social status.

In this study, a method of reconfiguring existing hemodialysis machines to perform hemodiafiltration, as hemodiafiltration has been shown to provide advantages over conventional hemodialysis, including various clinical advantages and overall improvement of patient survival.^[3]^ Older hemodialysis machines cannot be used for these purposes as they do not have any included functionality to provide fluid substitution to the blood. However, the proposed reconfiguration strives to make use of the existing functionalities of the hemodialysis machine to achieve this.

The proposed configuration uses an additional dialysis filter, termed the “replacement filter.” The primary purpose of this filter is to separate the dialysate inflow into two streams – one for dialysis, and the other for fluid substitution. There is a minor secondary purpose of this filter, which is to filter the dialysate prior to entering the hemodialysis filter which prevents any unnecessary contamination and fouling.

The hemodialysis machine used by Quantum Medical – the SURDIAL™ 55 – controls the inflow and outflow of the dialysate to maintain the ultrafiltration rate specified via user input. This proposed configuration takes advantage of this control to achieve more ultrafiltration than possible in a typical hemodialysis configuration. By partitioning a portion of the dialysate inflow for substitution, the actual dialysate outflow is lower than the dialysate outflow expected by the machine. As such, the dialysate is pumped out at a flow rate higher than anticipated, resulting in a higher actual ultrafiltration rate. While it is possible to increase the ultrafiltration rate manually in conventional hemodialysis, this may result in dehydration in the patient if increased excessively. With substitution of dialysate to the patient as found in hemodiafiltration, this would not be an issue.

In this research, the hemodialysis conditions will be replicated to match various key parameters provided by Quantum Medical. In the replicated set-up, human blood will be dialyzed in the conventional hemodialysis configuration, as well as in the proposed hemodiafiltration configuration. Using samples collected during the experiment, the urea levels will be monitored and compared to evaluate the efficiency of the proposed hemodiafiltration configuration as compared to that of the conventional hemodialysis configuration.

## MATERIALS AND METHODS

The dialysis set-ups were replicated with hemodialyzers (Elisio™ 17H) and tubing provided by Quantum Medical as well as various pumps and flow meters to control the flow. The tubing was configured to replicate the line set-up provided by Quantum Medical (Figure 1).

**Figure 1.**
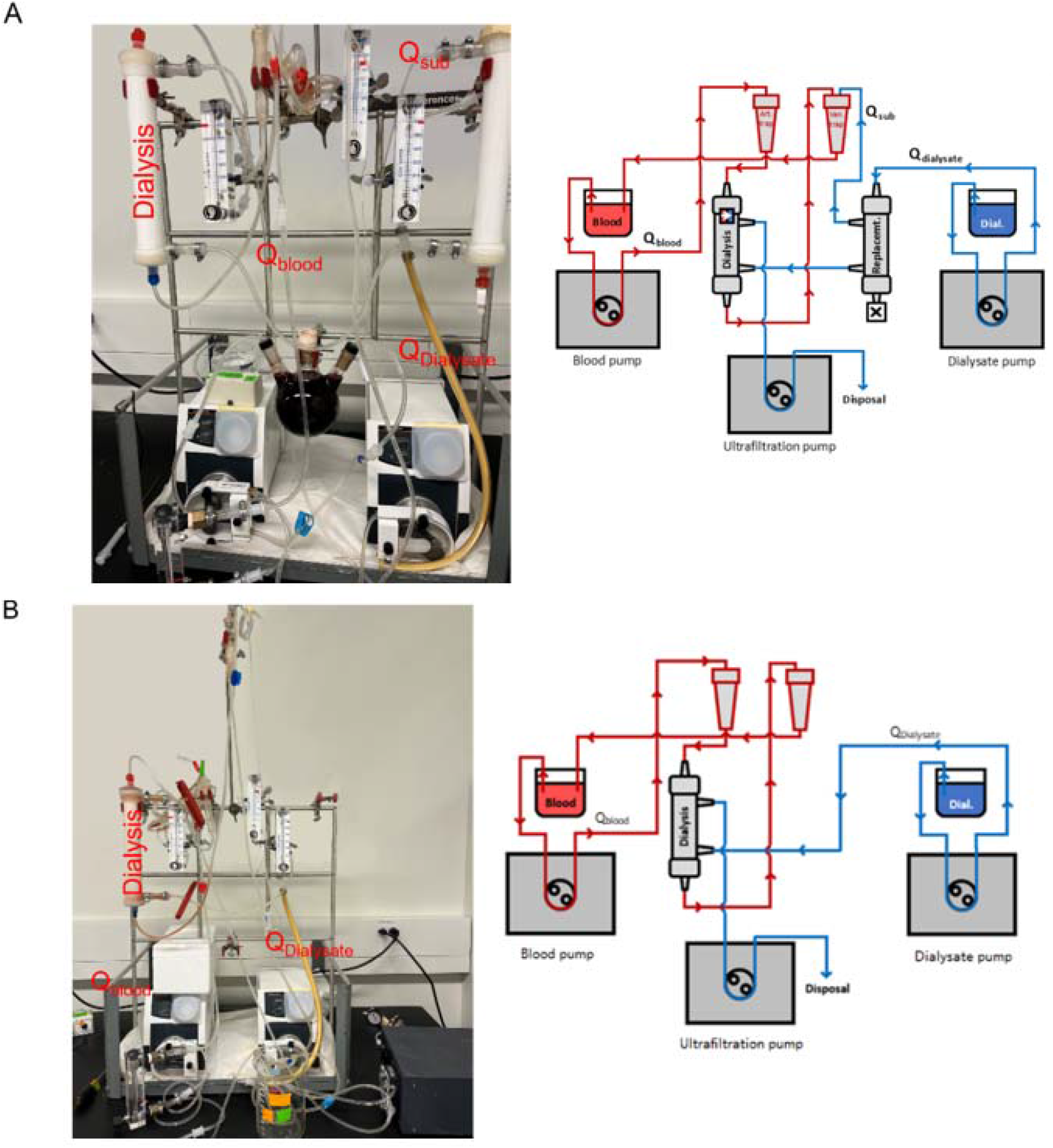
Hemodiafiltration (A) and hemodialysis (B) configurations.

Flow meters were placed in accordance with the locations of the flow sensors in the SURDIAL™ 55, the hemodialysis system used by Quantum Medical. Using these flow meters, the pump speeds were adjusted to match the following parameters (Table 1).

**Table 1.** Target parameters for dialysis.

| Parameter | Value |
| --- | --- |
| Dialysate flow rate | 600 mL/min |
| Blood flow rate | 400 mL/min |
| Substitution rate | 44 mL/min |
| Ultrafiltration rate | 8.3 mL/min |

### Blood Preparation

Citrated human whole blood was acquired (BioChemed Services) and spiked with urea to increase the concentration of urea by 30 millimolar. 500 milliliters of this urea-spiked human whole blood were then dialyzed against a citrate-based, bicarbonate-buffered dialysate for 18 minutes, with blood samples being collected at regular intervals.

### Assays

The efficiency of the machine was determined by comparing the urea clearance in hemodiafiltration and conventional hemodialysis.^[4]^ And human serum albumin and hemolysis were used to verify if there would be additional risk for the patient. During the dialysis, the plasma was extracted from the blood samples at intervals and analyzed for urea concentration via a urease/Berthelot reagent-based urea assay in Figure S1. The dialysate was also collected for analysis of human serum albumin via an enzyme-linked immunosorbent assay (Abcam) in Figure S2. Finally, the percent hemolysis of blood samples was also determined via spectrophotometry.

## RESULTS

### Urea Clearance

As the main goal of hemodialysis is to remove waste products and excess fluids from the blood of patients with kidney failure or chronic kidney disease who are unable to do so effectively on their own, it is important to quantify dialysis adequacy to improve the patient’s overall health and quality of life by preventing or reducing the symptoms and complications of kidney failure. The molecule urea was chosen as an indicator of dialysis adequacy as it is the most common marker for this purpose, owing to its ability to reflect biological changes in protein catabolism and kidney function, while also having an economical and straightforward assay.^[5-8]^ As shown in Figure 2, the urea level in the hemodiafiltration configuration decreases dramatically within 2 min as compared to conventional hemodialysis, which only achieves a low urea level at around 3 min. As shown in Table 2, the urea clearance was determined to be around 53.33±13.20 mL/min for the hemodialysis configuration and around 268.31±44.17 mL/min for the hemodiafiltration configuration. The urea reduction ratio (URR), another indicator of hemodialysis adequacy, is regarded as adequate above 65%.^[9-11]^ The URR was determined to be 76% for hemodiafiltration and 67% for hemodialysis. These data confirm that this hemodiafiltration set-up performs better than the hemodialysis set-up.

**Figure 2.**
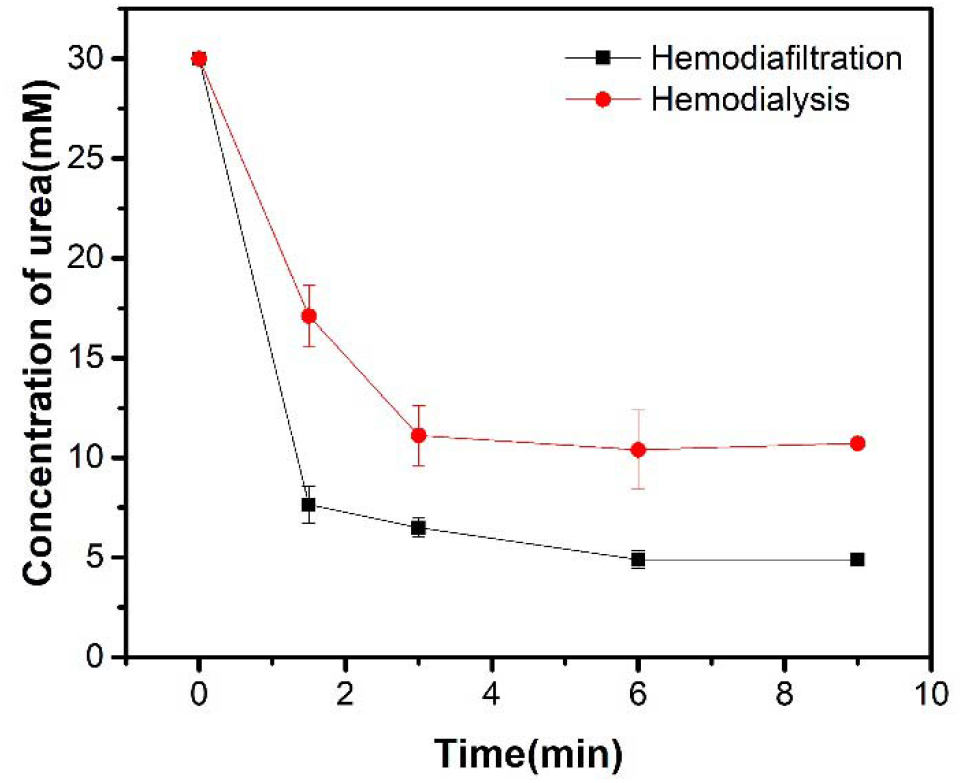
Decrease of urea concentration in blood sample with dialysis time.

**Table 2.** Summary of performance between hemodialysis and hemodiafiltration.

| Performance | Hemodialysis | Hemodiafiltration |
| --- | --- | --- |
| Urea Clearance | 53.33±13.20 mL/min | 268.31±44.17 mL/min |
| Albumin Loss | 0.23±0.025 µg/mL | 0.06±0.036 µg/mL |
| Hemolysis | 12.26±16.7% | 8.87±9.15% |

### Human Serum Albumin and Hemolysis

To further confirm that hemodiafiltration is as safe as the traditional hemodialysis approach, human serum albumin and hemolysis experiments were chosen as an indicator to verify the safety of the hemodiafiltration.^[12-14]^ As shown in Table 2, the human serum albumin measured from the dialysate was around 0.23±0.025 µg/mL in hemodialysis and around 0.06±0.036 µg/mL in hemodiafiltration. These values seem to be comparable and also negligible in comparison to the approximately 35 g/L found in the blood before dialysis. In the hemolysis experiment, the percent hemolysis increased by around 5.8±4.5% in the hemodialysis experiments and by around 5.5±3.49% in the hemodiafiltration experiments. There is no dramatic difference pre- and post-hemodiafiltration (Figure 3), but a slight increase in post-dialysis when comparing the blood pre- and post-hemodialysis. This may be due to conditions of the experiment such as the shear stress generated from the flow rates, different pressures, and patient-related medical conditions.^[15-17]^

**Figure 3.**
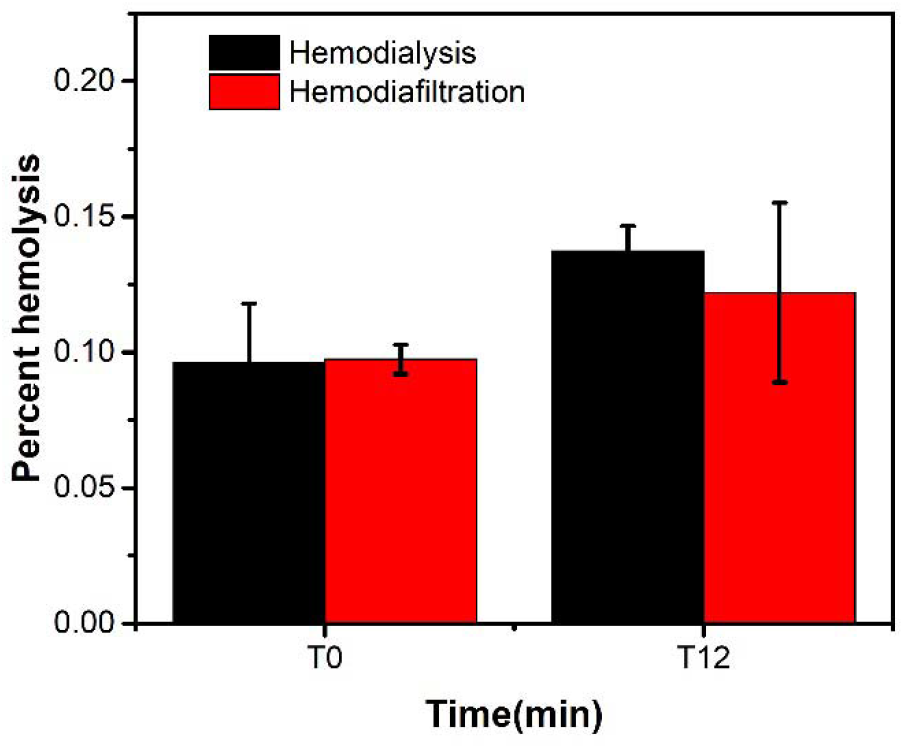
Percent hemolysis of the blood pre- and post-dialysis

## DISCUSSION

The increased urea clearance of the hemodiafiltration experiments suggests that the proposed hemodiafiltration configuration may indeed provide higher performance during each patient dialysis session. While the absolute K values may not be reflective of the exact values that are achieved *in situ*, the significant increase demonstrates that the hemodialysis is more efficient with the adaptation and provides a higher dose of dialysis.^[4]^ With the increased performance from the hemodiafiltration reconfiguration, patient serum urea concentrations should reduce more efficiently and thoroughly, which should afford the patient with the aforementioned clinical advantages and improved survival.

The amount of human serum albumin in the dialysate was measured to determine whether there would be any increased removal of albumin through the hemodialyzer. As the levels of human serum albumin in the hemodialysis experiment are similar to those in the hemodiafiltration experiment, there does not seem to be any significant effect of albumin loss due to the implementation of the proposed reconfiguration and thus does not increase the patient’s risk of hypoalbuminemia.^[18]^

The amount of hemolysis did not increase when the blood was dialyzed using the hemodiafiltration configuration when compared to that of the hemolysis configuration. This indicates that the different flow conditions of the proposed reconfiguration do not significantly increase the level of breakdown of blood cells during operation from its increased effective ultrafiltration and in this regard, does not pose any additional risk to the patient when used. It is worth noticing that there is obvious observable hemolysis pre-dialysis (Figure 3). However, this may be due to the condition of the blood during experimentation. As the blood used is not directly pumped from a patient as would be done in actual patient dialysis and has instead been drawn from a donor and stored for multiple days, the blood cells experience storage lesion and become more fragile and prone to hemolysis.^[15,19-20]^ In conjunction with the peristaltic pumps which may not be as optimized for pumping blood as those in a hemodialysis machine, a certain level of hemolysis is to be expected and may not reflect the level of hemolysis that may occur in the SURDIAL™ 55.

## CONCLUSION

In conclusion, the proposed hemodiafiltration reconfiguration appears to yield a better performance as compared to conventional hemodialysis when using the same parameters, such as blood and dialysate flow rate due to the increased ultrafiltration generated from the hemodialysis machine. The experimental measurements of human serum albumin and hemolysis do not indicate that the implementation of the proposed adaptation pose any additional risk to the patient in terms of albumin loss and increased *in vitro* hemolysis.

## ACKNOWLEDGMENTS

This work is supported by research funds from the Centre for Bioengineering and Biotechnology, University of Waterloo, and Quantum Medical.

## ETHICS STATEMENT

The study protocol involving purchased human blood was in accordance with the ethical standards and approved by the institutional review board.

