## Supplemental Data for "A Two-Filter Adaptation to Achieve Hemodiafiltration with Enhanced Performance"

| Urea (mM) | Absorbance |
| --- | --- |
| 2 | 0.996 |
| 1 | 0.54 |
| 0.5 | 0.231 |
| 0.25 | 0.117 |
| 0.125 | 0.053 |
| 0.063 | 0.026 |
| 0.031 | 0.012 |
| 0 | 0 |

Figure S1. Urea calibration curve from urease/Berthelot reagent-based urea assay.

| Human Serum Albumin (ng/ml) | Absorbance |
| --- | --- |
| 45 | 0.53525 |
| 22.5 | 0.3649 |
| 11.25 | 0.2661 |
| 5.63 | 0.20095 |
| 2.81 | 0.1803 |
| 1.41 | 0.1561 |
| 0.7 | 0.14605 |
| 0 | 0.1405 |

Figure S2. Human serum albumin calibration curve obtained using human serum albumin ELISA kit.
